# Alvis: a tool for contig and read ALignment VISualisation and chimera detection

**DOI:** 10.1101/663401

**Authors:** Samuel Martin, Richard M. Leggett

## Abstract

**Background:** The analysis of long reads or the assessment of assembly or target capture data often necessitates running alignments against reference genomes or gene sets. Aligner outputs are often parsed automatically by scripts, but many kinds of analysis can benefit from the understanding that can follow human inspection of individual alignments.

**Findings:** We developed Alvis, a simple command line tool that can generate visualisations for a number of common alignment analysis tasks. Alvis is a fast and portable tool that accepts input in the most common alignment formats and will output production ready vector images. Additionally, Alvis will highlight potentially chimeric reads or contigs, a common source of misassemblies. We found that splitting chimeric reads using the output provided by Alvis can improve the contiguity of assemblies, while maintaining correctness.

## Findings

### Background

Finding alignments between two sets of sequences is a fundamental task in bioinformatics. The analysis of long reads, the assessment of assembly results or evaluation of target capture protocols often necessitate alignment against reference genomes or gene sets. Many different tools exist to calculate alignments and these tools generate a range of different output formats. Most of the common formats consist of large tab separated lists which are designed for easy computer parsing rather than to convey intuitive human understanding. Yet many kinds of analysis can benefit from a visual depiction of alignments – for example, inspecting the layout of contigs across a chromosome, understanding where assemblies break down or picturing read coverage depth across a chromosome or gene. Such analysis can also expose the presence of chimeras formed by the artificial joining of disconnected parts of the genome.

To date, there are many tools available for alignment visualisation, which can vary in their specific function depending on the task the user faces. One possibility is to use a genome browser, such as ensembl[1] or IGV[2]. Genome browsers perform more than just sequence alignment visualisation, allowing the user to browse, search and analyse genomic sequence and annotation data. Genome browsers provide a graphical interface to users, many of which are available online through a web browser, which can be helpful for those less familiar with the command line. However, the wide range of tasks possible in a genome browser necessitates some complexity in the interface they provide and the images they produce are not usually of publication quality.

The tool Circos[3] is a popular tool for producing high quality images for publication, particularly for displaying many datasets which relate to a single genome (e.g. [4]), and for structural variation between genomes. However, while these diagrams look superficially attractive, they can lack clarity when overused and are typically of the wrong scale to visualise individual contig alignments.

The mummerplot tool, included in the MUMmer package[5], can create a dot plot of alignments between two sets of sequences, allowing easy assessment of rearrangements. These plots are simple, clear and easy to read, and the program itself has a simple command line interface which is easy to use. They do however require the user to use alignment software included in the MUMmer package, and can become diicult to read if there are many query or target sequences.

Another option is provided by BLAST’s web interface[6]. This allows the user to enter a sequence, which is queried against the NCBI nt database using BLAST. Alignments to the best hit are displayed on a diagram (similar to that in Figure 1a). This method, however, is unsuitable for analysing large numbers of sequences, or for comparing several alignments at once.

**Figure 1.**
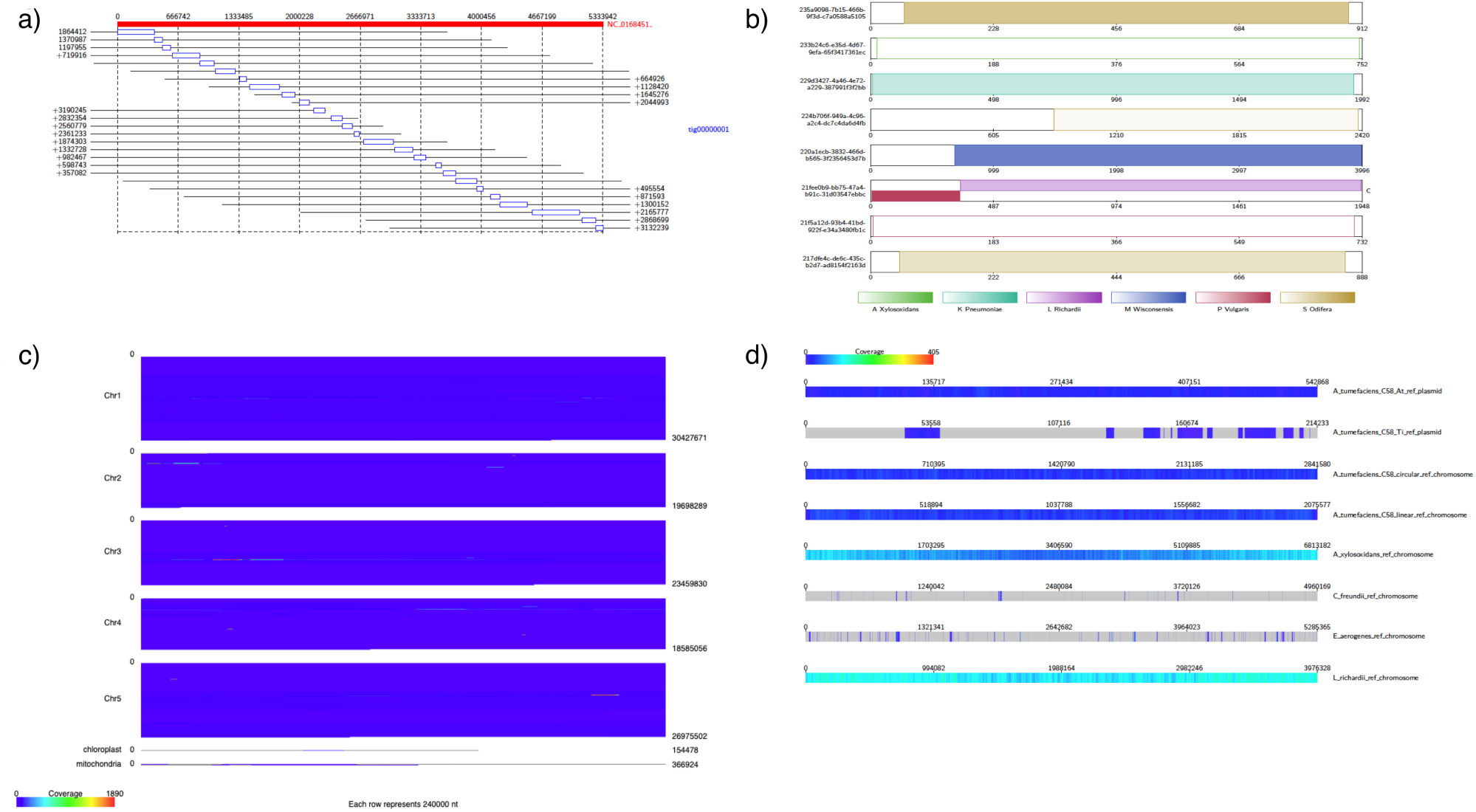
Example Alvis outputs: (a) Alignment diagram showing mapping of a contig against a reference genome. (b) Contig alignment diagram indicating a potential chimera in one of the reads (c) Genome coverage diagram showing reading coverages across *Arabidopsis thaliana* chromosomes. (d) Coverage map showing read coverage across a set of references.

Here we present Alvis, a tool for visualising alignments of long reads and assemblies which can generate four different types of publication quality diagram from a wide range of different input file types. Alvis enables flexible filtering of alignments by the user and will automatically highlight potentially chimeric reads or contigs. Furthermore, the user may choose to view only potentially chimeric sequences and obtain a text file containing a list of all such sequences and the approximate join location. We show that using this option on long read sequence data can improve the correctness and contiguity of assemblies.

### Description of Tool

Alvis is written in Java and can be run on any platform with a Java Runtime Environment e.g. Linux, MacOS and Windows. It has a simple command line interface for operation and runs rapidly. Alvis accepts inputs in the following formats: BLAST tabular, SAM files from BWA [7] and other aligners, PAF files from minimap2 [8], MUMmer’s .coords and .tiling files [5], PSL files from BLAT [9]. Additionally, the software has been designed to allow easy implementation of other formats. Diagrams can be output in either SVG or TeX formats. In the latter case, a LaTeX compiler is required to produce a PDF file.

The user can choose to filter alignments to remove noise. This is achieved by discarding alignments that are, by default, less than 1% of the query length (this value may be changed by the user). The user may also choose to display only alignments that could have come from chimeric reads. These are chosen when a query sequence contains exactly two non-overlapping alignments to either different target sequences or different sections of the same target sequence, that cover 90% of the query sequence. The length of each alignment must be at least 10% of the length of the query sequence. These default values may be adjusted by the user. Alvis can also output a separate plain text file containing the IDs of potentially chimeric reads or contigs, along with an approximate location of the join.

Alvis can be run from the command line using the Jar file provided in the distribution. For example, to create a coverage map from a minimap2 file:

`Java -jar Alvis.jar -inputfmt paf -outputfmt tex –type coverageMap -coverageType long -in alignments.paf -outdir output -out outprefix`

### Alvis Diagrams

#### Alignment Diagram

The alignment diagram (Fig. 1a) groups alignments by their target sequence, represented by a red bar at the top of the diagram. Each alignment is displayed as a rectangle underneath the bar, in line with the alignment position on the target. For a given target sequence, each alignment is ordered by the query ID and, for queries producing multiple alignments, by the start position of the alignment on the target. The length of each query sequence, and the position of the alignment on the query sequence, is indicated by a line parallel to the target bar and through the alignment rectangle. An example use might be to visualise the set of assembled contigs that align to a given chromosome.

#### Contig Alignment Diagram

The contig alignment diagram (Fig 1b) groups all alignments by their query sequence, and for each query, displays the ten longest alignments inside a rectangle representing the query sequence. These alignments are colour coded by target sequence and shaded to indicate position and orientation. With such a diagram, it is possible to view an assembled contig or long read to understand how discontinuously it maps to a target sequence and if it is chimeric.

#### Coverage Map Diagram

The coverage map diagram (Fig 1c) builds a coverage representation of a set of target sequences by dividing each into positional bins and incrementing the bin count for each query sequence that aligns to that bin. In the case of a query sequence with overlapping alignments, the largest alignment is chosen, and overlapping alignments are discarded. For each target sequence, a heat map image is produced with each bin represented by a pixel width. This image can be chosen to be either a square which wraps around, or a long bar (Fig 1c). Since each target sequence may have a different length, the pixel size is adjusted so that the heat map size is constant. An early version of Alvis was used to generate resistance gene coverage maps in [10].

#### Genome Coverage Diagram

The genome coverage diagram (Fig 1d) builds up a coverage representation in the same manner as the coverage map diagram. However, instead of producing individual target heatmaps, a single heatmap is produced which shows the coverage of every target sequence, in order, with a constant pixel size. This allows, for example, production of a single coverage map which shows the coverage of all chromosomes of a species by an assembly.

### Software Architecture

The software has been designed in a modular fashion to enable easy extension in the future. Central to the design are three abstract components; Alignments, Drawers, and Diagrams.

#### Alignments

Each alignment format is represented by a class implementing either the Alignment or DetailedAlignment interface, depending on the data that is available in that format. The DetailedAlignment class extends the Alignment class. Another class representing the alignment file parses the file and creates an array of Alignment or DetailedAlignment objects; this class must implement either the AlignmentFile interface, or the DetailedAlignmentFile interface (which extends the AlignmentFile interface). Thus implementation of a new alignment format requires just two short classes to be created.

#### Drawers

Similarly, the notion of an output format has been abstracted through the Drawer interface. Currently two classes implement the Drawer interface; one for TeX output, which uses the Tikz package, and one for SVG output. Further output formats can be implemented through writing a class which implements the Drawer interface. This interface contains methods for basic drawing operations, such as drawing a line between two points, and drawing text.

#### Diagrams

Alvis is capable of drawing four types of diagram. Each diagram takes an object implementing either the AlignmentFile or DetailedAlignmentFile interfaces, and creates in instance of a Drawer to output the diagram. The Alignment and Contig Alignment diagrams require an object implementing the DetailedAlignmentFile interface, while the Coverage Map and Genome Coverage diagrams require an object implementing either the AlignmentFile interface or the DetailedAlignmentFile interface.

## Methods

### Chimera Detection

To demonstrate the value of Alvis’ chimera detection function, we downloaded and assembled two sets of nanopore reads before and after chimera identification and removal. The first was sequenced from *Arabidopsis thaliana* accession KBS-Mac-74 [11] and the second was the Rel3 nanopore human dataset from the Nanopore WGS Consortium [12]. For each read set we mapped reads to the respective reference genome (TAIR10 [13]for *A. thaliana*, and GRCh38.p13 [14] for human) using min-imap2, creating two large paf files. These were parsed by Alvis to highlight suspected chimeric reads. Reads are identified as chimeric if they have exactly two distinct alignments to different regions of the reference genome, with both alignments together covering 95% of the read and each covering at least 10% of the read.

Using the “-printChimeras” option for Alvis produces a tab separated values file, where each line describes a potentially chimeric read, including the read ID, the approximate join position, and the two target sequences.

For the *A. thaliana* dataset we found a total of 2,817 out of 300,071 reads meeting Alvis’ chimera requirements. Of these, 264 reads had alignments to the same circular reference sequence (either mitochondria or chloroplast) and so were ignored, leaving 2,553 potentially chimeric reads. We then created a new read set, by copying all the original reads, and spitting each of the 2,553 potentially chimeric reads at the join position given by Alvis. A python script, chimera_splitter.py, is provided in the Alvis distribution to perform this automatically. Both the new read set and the original read set were assembled using flye[15]. A significant improvement in the contiguity of the assembly from the split reads is indicated in Table 1.

**Table 1.**
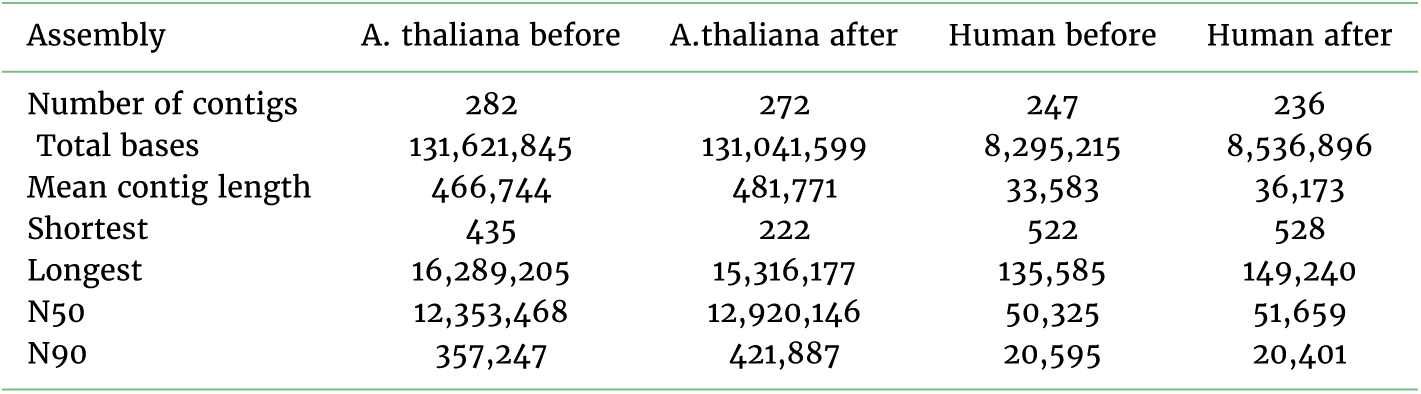
Contig statistics for flye assemblies of *A. thaliana* and human reads before and after chimera splitting.

| Assembly | A. thaliana before | A.thaliana after | Human before | Human after |
| --- | --- | --- | --- | --- |
| Number of contigs | 282 | 272 | 247 | 236 |
| Total bases | 131,621,845 | 131,041,599 | 8,295,215 | 8,536,896 |
| Mean contig length | 466,744 | 481,771 | 33,583 | 36,173 |
| Shortest | 435 | 222 | 522 | 528 |
| Longest | 16,289,205 | 15,316,177 | 135,585 | 149,240 |
| N50 | 12,353,468 | 12,920,146 | 50,325 | 51,659 |
| N90 | 357,247 | 421,887 | 20,595 | 20,401 |

For the human dataset, we found a total of 1,750 out of 658,224 chimeric reads, of which 90 had alignments to the mitochondrial sequence in the reference genome, so were ignored. This left 1,660 chimeric reads in total. As before, two assemblies were performed using flye; one from the original read set, and one from the read set where chimeras were split. The contiguity statistics for these assemblies are presented in Table 1.

## Summary

Alvis provides the ability to quickly and simply visualise contig and read alignments in four diagram types. A wide variety of popular file formats are supported and a simple API provides for future extension. Diagrams can be output as vector images, providing high quality publication-ready figures. The ability to output to SVG could allow future integration with web applications. Crucially, Alvis can also be used to highlight potential chimeric reads and contigs. In the former case, splitting chimeric reads can lead to a more accurate and contiguous assembly.

## Availability and requirements

**Project name:** Alvis

**Project home page:** https://github.com/SR-Martin/alvis

**Operating system:** Platform independent

**Programming language:** Java

**Other requirements:** Java Runtime Environment

**License:** GNU GPL v3

## Availability of supporting data

The data used to generate the diagrams in Figure 1 is available in the Alvis repository at https://github.com/SR-Martin/alvis. A detailed tutorial for creating these diagrams is available at https://alvis.readthedocs.io/en/latest/usage/example.html.

## Competing Interests

The authors declare that they have no competing interests.

## Funding

This work was strategically funded by the BBSRC, Core Strategic Programme Grant BB/CSP17270/1 at the Earlham Institute.

